# Developmental Progression and Single-Cell Heterogeneity of *Orientia tsutsugamushi* during Intracellular Infection

**DOI:** 10.64898/2025.12.09.693123

**Authors:** Chitrasak Kullapanich, Frances Aylward, Rachel Jackson, Jeanne Salje

## Abstract

*Orientia tsutsugamushi*, the causative agent of human scrub typhus disease, is an obligate intracellular bacterium transmitted by mites. It replicates exclusively within eukaryotic cells, inhabiting a range of host cell types in mammals and mites. Previously, we demonstrated that its extracellular, non-replicative form is developmentally distinct from the intracellular replicative form. In this study, we investigated spatial and temporal heterogeneity within the intracellular bacterial population. By quantifying bacterial DNA replication, protein synthesis, cell division, and subcellular localization, we defined four distinct intracellular growth stages: pre-active, lag, log, and maturation. Using microscopy-based single-cell analysis of selected transcripts and proteins, we further revealed transcriptional and translational heterogeneity within microcolonies. These findings highlight the complex, asynchronous nature of intracellular *O. tsutsugamushi* growth and underscore the importance of single-cell analysis in understanding its pathobiology.

**Importance:** *Scrub typhus* is a serious, sometimes life-threatening disease caused by the bacterium *Orientia tsutsugamushi*, which is spread by the bites of tiny mites and is common in many rural parts of Asia. This bacterium can only survive and grow inside living cells, where hundreds of individual bacteria may infect a single host cell. Until now, it was unclear whether all the bacteria inside a cell behaved the same way or if they existed in different stages of growth. In this study, we show that *O. tsutsugamushi* goes through several distinct stages while growing inside host cells and that not all bacteria are at the same stage at the same time. Even within a single infected cell, some bacteria are actively dividing, while others are preparing to leave the cell. Understanding these stages provides a new framework for exploring how the bacterium develops, spreads, and causes disease — insights that could help guide future research and improve ways to control or treat scrub typhus.

## Introduction

*Orientia tsutsugamushi* (Ot) is an obligate intracellular bacterium and the causative agent of scrub typhus, a mite-borne febrile illness endemic across much of rural Asia and the Pacific region^1^. Scrub typhus poses a significant public health burden, often representing one of the most common causes of acute undifferentiated fever in endemic areas^2,3^. Transmission occurs through the bite of infected larval mites (chiggers), in which Ot is maintained via transovarial inheritance^4^. Humans, however, are accidental hosts and represent a dead-end for the bacterial infection. Once in the human body, Ot disseminates via the lymphatic and circulatory systems, establishing infection in a wide range of tissues, including the liver, lungs, spleen, and central nervous system^5^. If left untreated, scrub typhus can result in multi-organ failure and death, with an estimated median mortality rate of 6% in untreated cases^6^.

The intracellular lifecycle of Ot has been characterized morphologically and at the level of host-pathogen interactions, but less is known about the bacterium’s own developmental changes during the course of a multi-day infection within host cells. Ot enters host cells through clathrin-mediated endocytosis^7^ and macropinocytosis^8^, subsequently escaping the endolysosomal pathway to reside freely in the host cytoplasm^7^. Once cytosolic, the bacterium manipulates the host environment to promote survival and replication. Ot actively modulates autophagy, both inducing and evading this pathway to suit its needs^9,10^. It hijacks the host microtubule network to traffic within the cytoplasm^11^, a process mediated by the bacterial outer membrane protein ScaC^12^. Replication occurs in densely packed microcolonies typically localized near the host nucleus, although this positioning can vary depending on the bacterial strain^13^. Upon completion of replication, Ot transitions to the cell periphery and exits the host via a budding process that retains host plasma membrane^14^. Notably, these extracellular, budded bacteria exhibit distinct morphological and molecular features, suggesting a developmentally specialized, non-replicative state^8^.

Despite these advances in understanding host cell manipulation and bacterial replication, the intracellular developmental program of Ot itself remains poorly defined. The question of whether Ot undergoes discrete, stage-specific changes while inside host cells has not been thoroughly addressed. Moreover, it is not known whether individual bacterial cells within a microcolony behave homogeneously or whether they exhibit functional heterogeneity, as has been observed in other bacterial systems^15^.

Single-cell heterogeneity has emerged as a critical aspect of bacterial pathophysiology in various species^16^. In biofilms, for example, genetically identical cells differentiate into distinct subpopulations with specialized roles, such as matrix production, motility, and dormancy^17,18^. Similar cell-to-cell variability is observed in sporulation in *Bacillus subtilis*, where only some cells enter the sporulation pathway^19^; in persistence in *Escherichia coli*, where rare dormant cells tolerate antibiotics^20^; in competence development in *Streptococcus pneumoniae*^21^; in type VI secretion system activation in *Vibrio cholerae*^22^; in intracellular fate decisions during *Salmonella enterica* infection^23^; and in differential phage susceptibility outcomes in *E. coli* populations^24^. These examples underscore how phenotypic heterogeneity, even among clonal bacterial populations, can confer survival advantages in fluctuating environments or host niches.

In this study, we sought to investigate whether Ot exhibits similar single-cell heterogeneity during its intracellular lifecycle. We first defined four broad developmental stages of intracellular infection—pre-active, lag, log, and maturation—based on DNA replication, cell division, protein synthesis, and position of bacteria. Focusing on the log phase, we then used microscopy-based quantification of transcript and protein levels in individual bacterial cells to assess variability within microcolonies. Our findings reveal that even within a single host cell, *Ot* populations are not homogeneous but instead comprise distinct subpopulations with differential gene expression patterns. These results provide new insight into the developmental complexity of *O. tsutsugamushi* and establish a foundation for understanding how phenotypic diversity contributes to its intracellular survival and pathogenesis.

## Results

### Bacteria progress through four stages of growth during intracellular infection

We sought to define the distinct stages of bacterial growth during intracellular infection by Ot. Bacteria outside eukaryotic host cells, prior to infection, are mostly metabolically inactive due to a lack of nutrient availability. It has been shown that these extracellular bacteria do not exhibit protein synthesis activity based on incorporation of a clickable methionine amino acid analog, L-homopropargylglycine (HPG) that is conjugated to a fluorescent probe after sample fixation^8^. We used the same microscopy-based amino acid incorporation assay to ask how soon after infection of murine fibroblast L929 cells bacteria initiate detectable protein synthesis activity. We observed measurable activity in 30% of bacteria at 1.5 hours post infection, the earliest time point it is possible to collect (Fig. 1A, B). This did not increase significantly between 1.5- and 7.5-hours post infection both in terms of the amount of activity, measured by fluorescence intensity (Fig. 1A), and the percentage of bacteria exhibiting measurable signal (Fig. 1B). Both measures exhibited a significant increase at 25.5 hours post infection. Consistent with previous reports, bacteria that had not infected cells and were located outside host cells exhibited no measurable incorporation of HPG (red spots in Fig. 1A). This led us to conclude that at least some intracellular Ot cells become at least somewhat metabolically active as early as 1.5 hours post infection. By contrast, we define those bacteria that are not yet measurably active in protein synthesis as pre-active.

**Figure 1.**
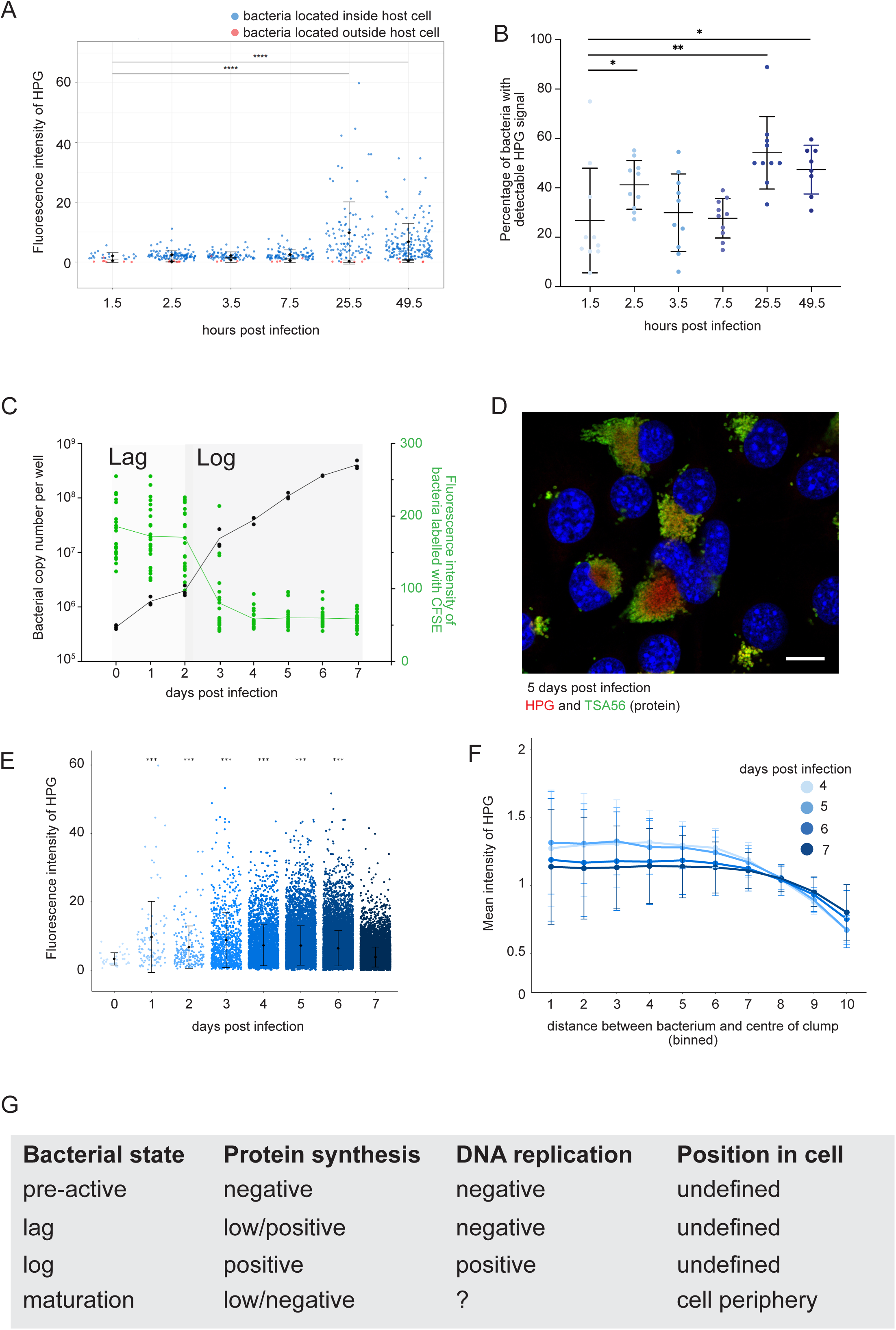
Defining different stages of Ot over time. A. Graph showing the fluorescence intensity of individual bacteria labelled with the protein synthesis reporter probe HPG at early time points after infection. Bacteria were classified as being located inside the host cell or outside the host cell based on microscopy, and these are represented in blue and red respectively. A Kruskal-Wallis test was used to determine the significance at each time point compared to the earliest time of 1.5 hours post infection. **** P ≤ 0.0001. The following minimum number of bacteria were counted from across three biological replicates: HPG IB ≥ 21, HPG EB ≥4, TSA56 IB ≥86, TSA56 EB ≥ 4. B. Graph showing the percentage of bacteria with detectable HPG signal at different times after infection. At least 51 infected cells were counted from across three biological replicates. Statistical significance was determined using Mann-Whitney test. * p ≥ 0.05, ** p ≥ 0.01. C. Graph showing the fluorescence intensity of individual live bacterial cells labelled with the stable dye CFSE. A decrease in CFSE intensity results from bacterial division. The bacterial genome copy number is measured using qPCR against the single copy gene *tsa47* and this is plotted on the same graph. Bacterial genome copy number increases over time as CFSE intensity decreases through cell division. D. Representative image of Ot in L929 cells labelled with HPG (red) and an antibody against the surface protein TSA56 (green). Nuclei are labelled with Hoechst in blue. Scale bar = 10 µm. E. Graph showing the fluorescence intensity of individual bacteria labelled with HPG at 1-7 days post infection. A Kruskal-Wallis test was used to determine the significance at each time point compared to the earliest time point. *** P ≤ 0.001. F. Graph showing the fluorescence intensity of individual bacteria labelled with HPG at different positions within an infected host cell. The fluorescence intensity relative to bacterial location is shown in separate curves at 4-, 5-, 6- and 7-days post infection. There is a decrease in fluorescence intensity in bacteria located towards the edge of an infected cells, consistent with the image shown in D. G. Summary of the four distinct stages of bacterial growth based on the data presented in this figure.

Next, we asked how soon after infection bacteria begin exponential growth and whether this was preceded by a lag phase as seen in other bacteria. We measured both DNA replication and cell division, because it is possible that bacteria could undergo cell division prior to DNA replication if multiple genome copies were present at the time of infection. To measure DNA replication we determined the bacterial genome copy number at each time point using quantitative PCR. To measure cell division we quantified fluorescence intensity of a stable fluorescent probe, CFSE, which was loaded into bacteria prior to infection of host cells. This probe has been shown to be stable over several days in Ot in the absence of division^25^, and the intensity decreases upon cell division due to dilution into daughter cells. We applied both analyses to the same sample of bacteria grown in L929 cells and observed that both measurements indicate a lag phase of slow growth, shown by low levels of both DNA replication and cell division, during the first two days of infection (Fig. 1C). This was following by initiation of cell division between 2- and 3-days post infection, with higher rates of DNA replication continuing up to 7 days post infection. Note that CFSE dilution can only be used to assess the onset and early stages of cell division because after a few rounds of cell division it has been diluted to undetectable levels. These data demonstrate that cell division and DNA replication are coupled in Ot during early stages of growth and confirms that intracellular Ot undergoes lag and log phases of growth akin to other bacteria.

We asked whether Ot undergoes a final stage of growth in which it prepares for exit from host cells. This would be similar to stationary phase growth in free living bacteria, which occurs when nutrient levels are depleted. Since this stage may occur concurrently with log phase growth in infected cells, it is not possible to define it by the bulk measurements of bacterial copy number using qPCR. CFSE dilution analysis is also not a tool that cannot be used to measure late stages of growth. Therefore we turned to two different measures to define this late-stage population. First, we measured HPG incorporation using fluorescence microscopy to assess whether there was a decrease in protein synthesis activity. Second, we used the subcellular position of bacteria to define different bacterial populations during late stages of infection reasoning that bacteria preparing to exit infected cells by budding would be located at the cell periphery. We measured HPG levels and bacterial subcellular position to analyse late stages of growth in an infected cell. We observed bacteria at the periphery of infected cells at late stages of infection, and these had a lower average HPG intensity than those adjacent to the nucleus (Fig. 1D-F). This led us to define a final stage of growth in the intracellular cycle of Ot, the maturation stage, in which bacteria are located on the surface of infected cells prior to egress as extracellular bacteria.

Together, these assays lead to the identification of four distinct stages of growth in intracellular Ot: pre-active, lag, log and maturation stages. It is expected that individual Ot cells will transition through these sequentially as they grow and divide. These stages are summarized in Fig. 1G.

### The timing of transition to the maturation stage depends on bacterial copy number within an infected cell

Maturation phase is the final growth stage in the intracellular lifecycle of Ot that appears as bacteria are preparing to exit from infected host cells. It was unclear whether transition to this stage of growth was dependent on the chronological time following infection, or the number of bacteria in an infected cell. To address this question, we quantified the number of Ot in the maturation phase of growth at different times after infection and compared this between cells that had been infected with different initial levels of bacteria (Fig. 2A). We carried out this experiment in human umbilical cord endothelial cells (HUVECs), because their large cytoplasm enabled clear separation between bacteria located in the perinuclear colony (log phase) and bacteria located at the edge of infected cells (maturation phase). We observed that individual HUVEC cells could classified into three groups based on the positions of bacteria within the cell. These groups are: 1. cells with bacteria primarily clustered in the perinuclear region, 2. Cells with bacteria in a perinuclear cluster and dispersed through the cytoplasm and 3. Cells with bacteria in a perinuclear cluster, dispersed through the cytoplasm, and located at the cell periphery. We reason that these three groups are progressively transitioning to a state in which bacteria are actively exiting the infected cell. We quantified the proportion of cells in each population at different points in time following infection with different bacterial doses (Fig. 1B).

**Figure 2.**
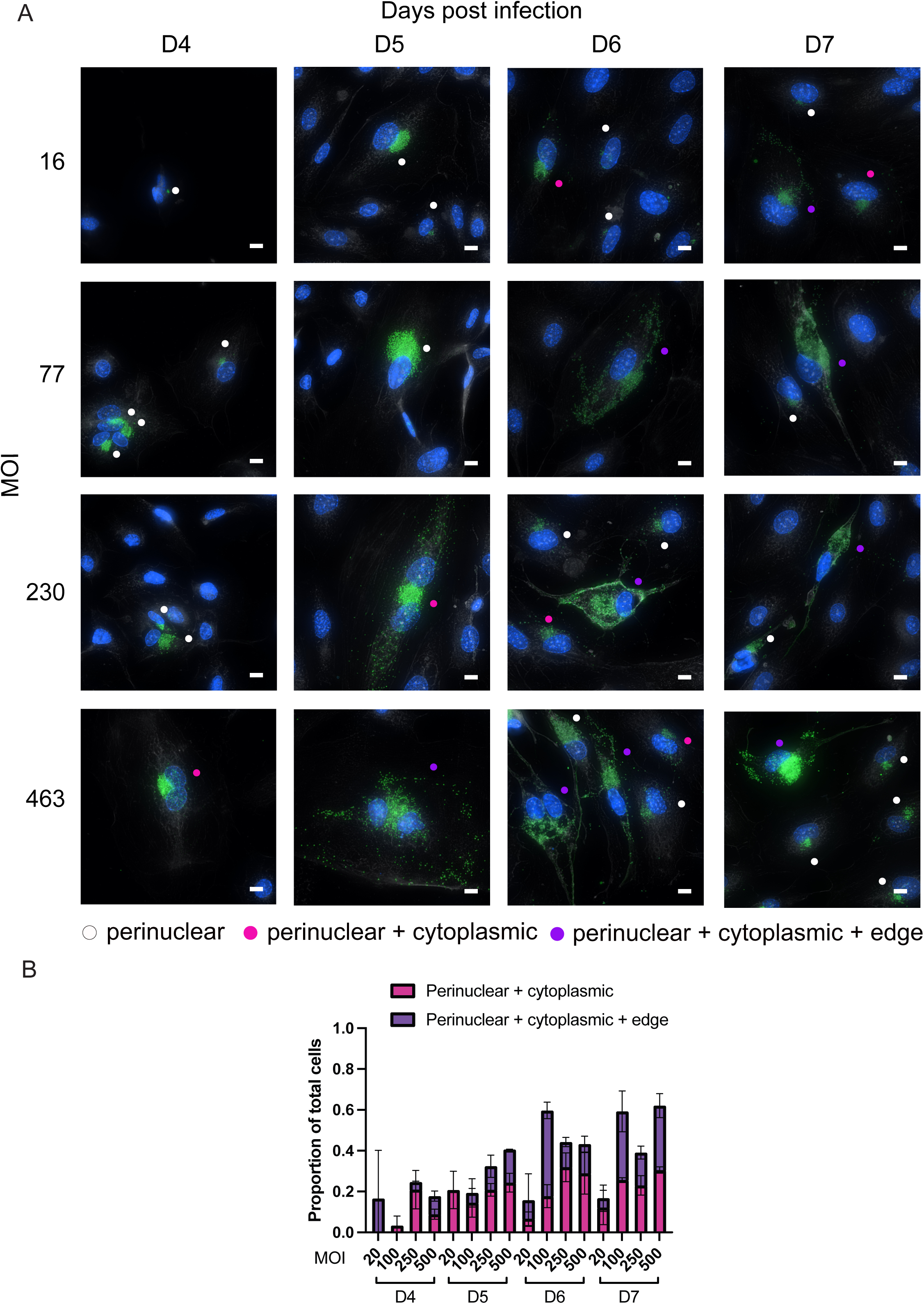
The timing of bacterial maturation is dependent on multiplicity of infection. A. Representative immunofluorescence microscopy images of HUVEC cells infected with Ot at MOIs of 16, 77, 230 and 463 and fixed at 4-, 5-, 6-, and 7-days post infection. Bacteria are labelled in green using an anti-TSA56 antibody, host nuclei are labelled with Hoechst and shown in blue. Infected cells are scored as having one of three distributions of bacteria: perinuclear; perinuclear + cytoplasmic; perinuclear + cytoplasmic + edge. These scores are shown by circles within images, colored in white, pink and purple respectively. B. Quantification of the number of host cells scored into the three classes of bacterial distribution shown in A. These data are taken from >1000 cells across two biological replicates. Both timepoint and MOI significantly affected proportion of maturation cells (binomial GLM, p<0.001). The Timepoint x MOI interaction was also significant (p<0.001), so effects were not interpreted in isolation.

The proportion of bacteria in group 3, in which the cell is primed for bacterial exit, is low at 4-days post infection, but increases substantially at 5- and 6-days following infection. The increase in proportion of these cells primed for exit was dependent on the starting MOI, with higher bacterial doses leading to earlier increases in cells with bacteria located at the cell periphery. This demonstrates that transition to this growth stage is dependent on bacterial number rather than purely chronological time. Notably, a decrease in cells containing bacteria at the periphery was observed at 7-days post infection in all but the lowest MOI conditions. This indicates that bacterial egress may be a finite process in infected cells, leaving a residual perinuclear microcolony at late stages of infection.

### There is heterogeneity between individual bacterial cells within an intracellular microcolony

During log phase growth, roughly 2-7 days post infection in our experimental set up, bacteria replicate in a tightly packed microcolony located adjacent to the eukaryotic host cell nucleus. Given that this is followed by a distinct stage of growth in which bacteria move to the edge of the cell in preparation for exit, and given that extracellular bacteria are known to be in a distinct developmental stage, we asked whether there was population heterogeneity within the bacterial microcolony during log phase growth, which might underpin the selection of individual bacteria that transition to the subsequent maturation and extracellular forms of growth. To address this question we labelled bacteria with antibodies or RNAscope probes against seven (antibodies) or eight (RNAscope) genes and determined the patterns of protein expression or transcript levels in individual bacteria within a microcolony. This experiment was carried out in L929 cells and representative images showing protein and transcript levels of the transcriptional regulator CtrA and the surface protein ScaA are shown in Fig. 3A. First, we analysed the total fluorescence intensity of each protein and transcript at different times post-infection (Fig. 3B-C). Whilst there was a general increase after infection, the patterns of gene expression differ between different genes. CtrA, TrcR and RpoD are all proteins involved in the regulation of gene expression. CtrA exhibited the earliest increase in expression at both the transcript and protein level, consistent with its known role in other bacteria as a master regulation of gene expression^26,27^. FtsZ and FtsQ are both members of the bacterial divisome^28,29^. FtsZ is involved in bacterial septation and the levels of transcript and protein both increase over time, with a trend that matches bacterial growth as determined by bacterial DNA copy number (Fig. 1A). By contrast, the patterns of the peptidoglycan synthesis regulator FtsQ are different than FtsZ with protein levels peaking at days 3-4 and then reducing at later times in infection. ScaA and ScaC are both bacterial surface proteins. ScaC drives bacteria trafficking along microtubules^12^ and the role of ScaA is unknown. ScaA protein and transcript levels are generally higher than ScaC, and transcript levels increase during infection. This is consistent with high levels of *scaA* RNA in extracellular bacteria reported previously^8^. Taken together, these data show that different genes are expressed differentially over time during the intracellular infection cycle of Ot.

**Figure 3.**
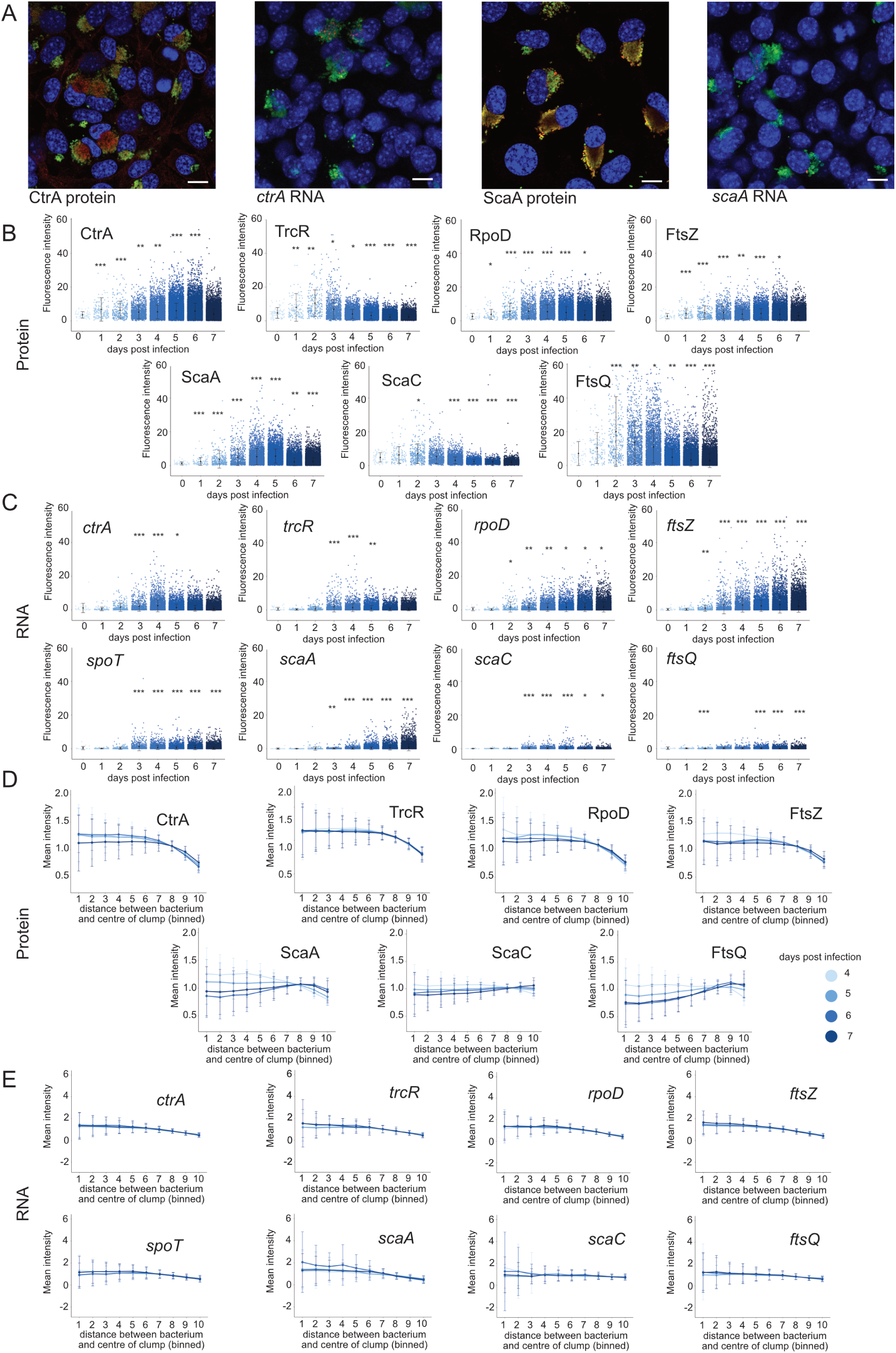
Heterogeneous gene expression within bacterial microcolonies. A. Representative images of immunofluorescence microscopy and RNAscope microscopy of Ot infected in L929 cells fixed at 5 days post infection, showing localisation of bacteria expressing *ctrA* and *scaA*. Immunofluorescence microscopy labels proteins using a polyclonal antibody against the protein of interest (CtrA, ScaA) in red. The bacteria are counterstained using an antibody against TSA56 in green and host nuclei are labelled with Hoechst in blue. Scale bar = 10 µm. B. Quantification of the fluorescence intensity of individual bacteria labelled with antibodies against seven different Ot proteins at 0-7 days post infection. All images with each antibody were acquired using constant laser intensity and imaging settings. C. Quantification of individual bacteria labelled with probes against transcripts of eight Ot genes. All images with each probe were acquired using constant laser intensity and imaging settings. Statistical significance in C and D were determined using a Kruskal-Wallis test. * p ≤ 0.05, ** p ≤ 0.01, *** p ≤ 0.001. D. Graph showing the fluorescence intensity of individual bacteria labelled with seven different antibodies at different positions relative to the center of a bacterial microcolony within an infected host cell. The fluorescence intensity relative to bacterial location is shown in separate curves at 4-, 5-, 6- and 7-days post infection. E. Graph showing the fluorescence intensity of individual bacteria labelled with eight different RNAscope probes at different positions relative to the center of a bacterial microcolony within an infected host cell. The fluorescence intensity relative to bacterial location is shown in separate curves at 4-, 5-, 6- and 7-days post infection.

We then asked whether the levels of these proteins and transcripts varied between bacteria located in different positions of the cell at a single point in time. Figs 3D and E show the fluorescence intensity of bacteria across a range of distances away from the center of the perinuclear microcolony, measured at days 4-7 post infection. We observed two general patterns of protein expression. In the first, the protein is more abundant in bacteria in the middle of the microcolony, i.e. closer to the nucleus. This pattern in observed in CtrA, TrcA, RpoD, FtsZ at all timepoints and on ScaA at days 3 and 4 post infection (Fig. 3A, 3D). By contrast, ScaC, FtsQ at all timepoints and ScaA on days 4 and 5 post infection exhibit a different pattern of expression whereby protein levels are higher on bacterial cells located further away from the centre of the microcolony (Fig. 3A, 3D). The patterns of genes with decreasing protein expression towards the outside of the microcolony were consistent at the transcript level (Fig. 3E). *ctrA, trcR, rpoD* (all time points) and *scaA* (early time points) exhibit a decrease in bacteria further from the nucleus. By contrast *ftsQ, scaC* (all time points) and *scaA* (late time points) and *spoT,* for which an effective antibody was not available, exhibit only a slight decrease or no change between bacteria in the centre and at the edge of the microcolony (Fig. 3E). Inspection of the images reveals that in these cases the bacteria expressing high levels of these transcripts are present at low numbers throughout the microcolony (Fig. 3A).

We sought to further explore the different patterns of transcript levels within a bacterial microcolony. We selected six genes, *ctrA, rpoD rpoH, scaA, spoT* and *tsa56,* and carried out low throughput, high quality microscopy imaging to compare the overall patterns of expression within a bacterial microcolony in mouse fibroblast L929 cells and human umbilical vein endothelial HUVEC cells at 5 days post infection (Fig 4). These data demonstrate very high transcript levels for *tsa56,* consistent with RNA sequencing data^30^. The other genes are present in much lower copies per cell, and not present in all bacteria. Where present, transcripts are present in single puncta which may correspond to individual transcripts. These data show that the transcripts exhibit broadly similar patterns of distribution across at least two different cell types.

**Figure 4.**
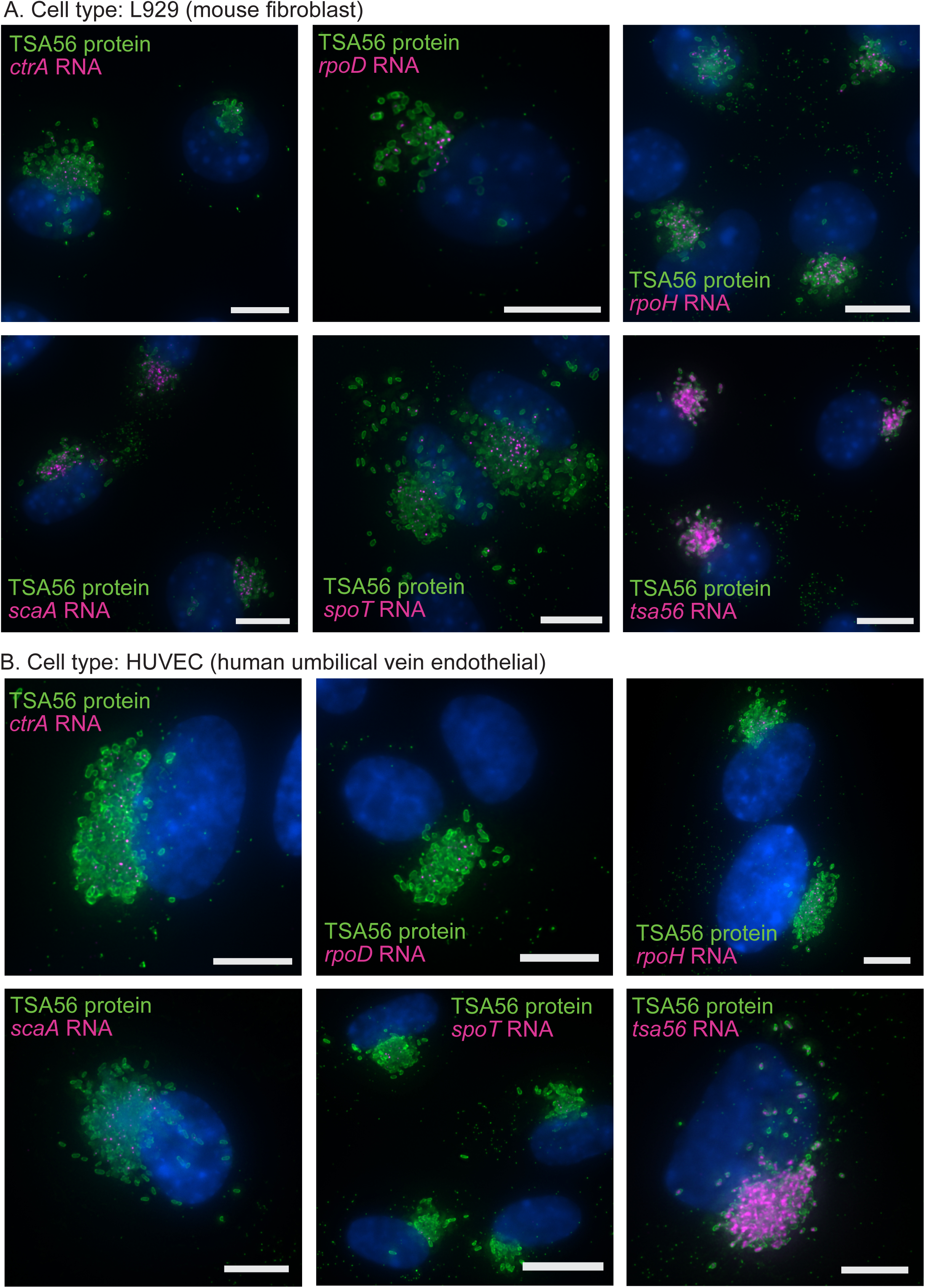
Different cell types have similar patterns of transcript distribution as determined by RNAscope. Representative images of Ot bacteria grown in L929 (A) or HUVEC (B) and labelled with probes against six different bacterial transcripts (*ctrA, rpoD, rpoH, scaA, spoT* and *tsa56*). Transcripts are shown in magenta and bacteria are counterlabelled with an antibody against TSA56 (green). Host nuclei are labelled with Hoechst and shown in blue. Scale bar = 10 µm.

### The intracellular infection cycle of cells infected with intracellular bacteria and extracellular bacteria are similar

It is known that the extracellular stage of Ot is a distinct developmental stage compared with the intracellular stage, with different shapes and protein profiles^8^. Both are known to be infectious. All experiments shown up to this point were carried out using infection with intracellular bacteria that had been mechanically isolated from infected cells. We asked whether the stages of growth defined above, and the spatial heterogeneity in gene expression, would differ if the infection was initiated using by extracellular bacteria that had previously budded off the surface of infected cells.

A comparison of the timing of initiation of protein synthesis, as measured by HPG incorporation, between L929 cells infected with IB-form Ot and EB-form Ot revealed that the overall timing and intensity of HPG incorporation was similar between the two (Fig. 5A), although there was significantly higher incorporation in IB infected bacteria at 25.5 hours post infection. This difference was gone by 49.5 hours post infection and indicates an earlier activation of bacteria from IB form.

**Figure 5.**
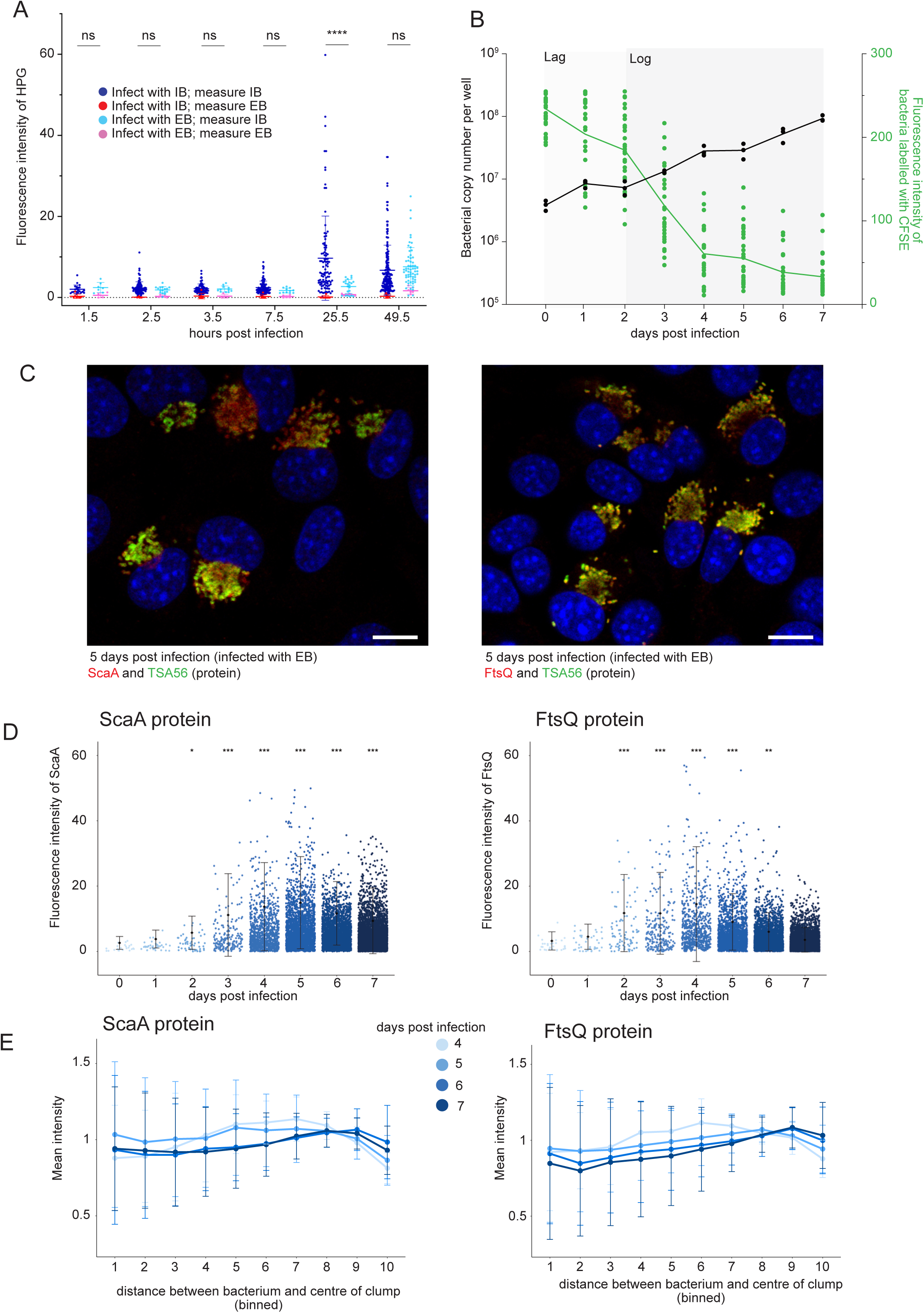
Analysis of bacterial growth following infection with EB Ot. A. Graph showing the fluorescence intensity of individual bacteria labelled with the protein synthesis reporter probe HPG at early time points after infection. Cells were infected with IB Ot or EB Ot, and then bacteria in fixed cells at different times after infection were classified as being located inside the host cell (IB) or outside the host cell (EB) based on microscopy. A Kruskal-Wallis test was used to determine the significance at each time point compared to the earliest time of 1.5 hours post infection. Ns = not significant, **** P ≤ 0.0001. The minimum number of bacteria counted from across three biological replicates was as follows: HPG IB ≥7, HPG EB ≥ 4, TSA56 ≥ 20, TSA56 ≥ 4. B. Graph showing the fluorescence intensity of individual live bacterial cells labelled with the stable dye CFSE. A decrease in CFSE intensity results from bacterial division. The bacterial genome copy number is measured using qPCR against the single copy gene *tsa47* and this is plotted on the same graph. Bacterial genome copy number increases over time as CFSE intensity decreases through cell division. C. Representative images of immunofluorescence microscopy of L929 cells infected with EB Ot and fixed at 5 days post infection, showing localisation of ScaA and FtsQ. Immunofluorescence microscopy labels proteins using a polyclonal antibody against the protein of interest (ScaA, FtsQ) in red. The bacteria are counterstained using an antibody against TSA56 in green and host nuclei are labelled with Hoechst in blue. Scale bar = 10 µm. D. Quantification of the fluorescence intensity of individual bacteria labelled with antibodies against ScaA and FtsQ at 0-7 days post infection with EB Ot. All images with each antibody were acquired using constant laser intensity and imaging settings. Statistical significance was determined using a Kruskal-Wallis test. * p ≤ 0.05, ** p ≤ 0.01, *** p ≤ 0.001. E. Graph showing the fluorescence intensity of individual bacteria labelled with antibodies against ScaA and FtsQ at different positions within a host cell infected with EB Ot. The fluorescence intensity relative to bacterial location is shown in separate curves at 4-, 5-, 6- and 7-days post infection.

**Figure 6.**
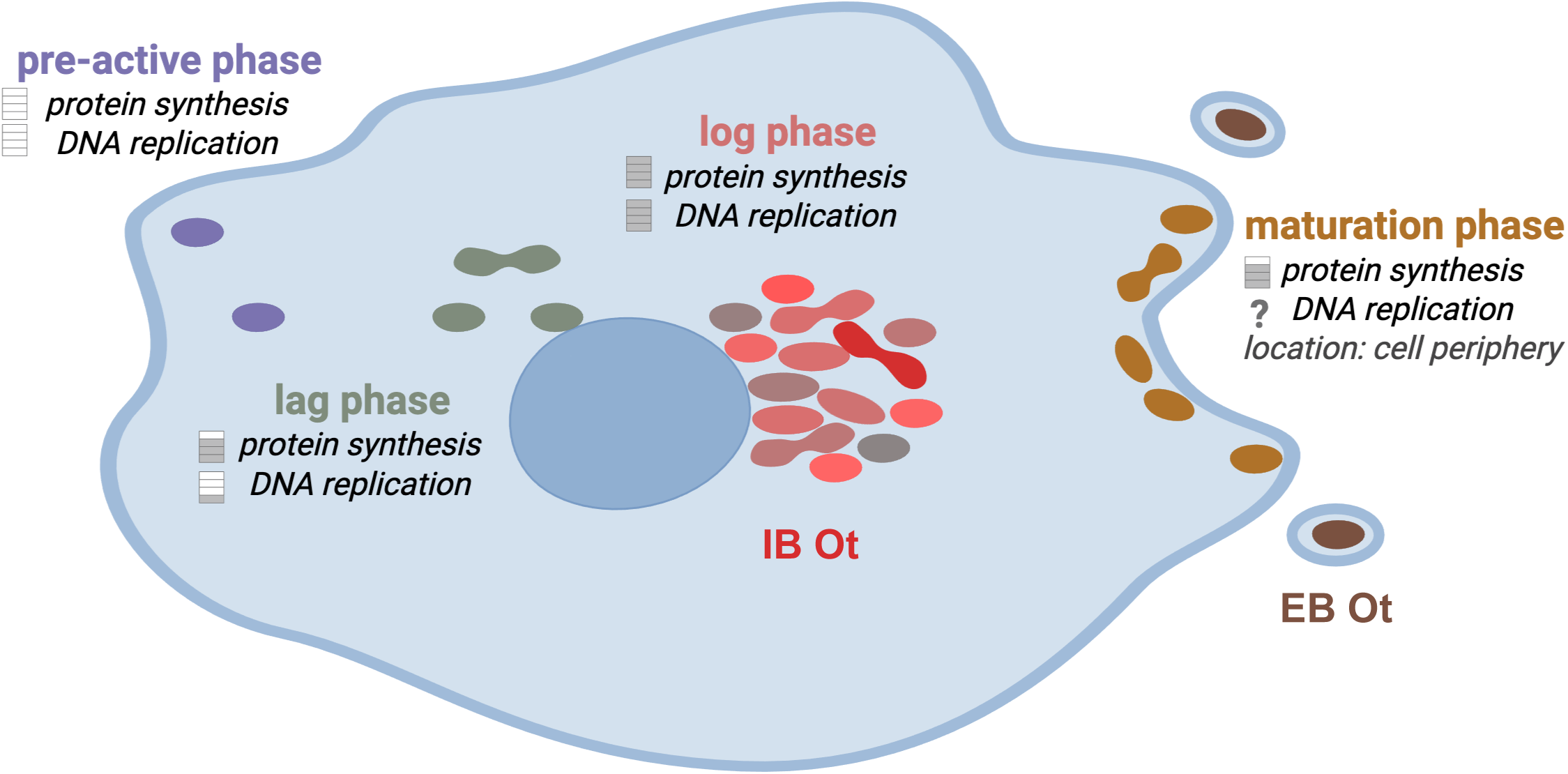
Overview of the stages of intracellular growth in Ot infection. Schematic diagram summarizing the distinct stages of growth during intracellular growth of Ot. The relative levels of protein synthesis activity and DNA replication activity at different stages of growth are shown. IB Ot and EB Ot are shown inside and outside an infected cell respectively.

We measured the timing of cell division and DNA replication in EB-form Ot following infection of L929 cells (Fig. 5B). Similar to IB infections, the log phase of growth began at 24 hours post infection. However there was less of a sharp difference between the two phases, and overall DNA replication was lower than for IB.

Finally, we selected two proteins FtsQ and ScaA, and measured their timing and distribution in L929 cells infected with EB-Ot (Fig. 5C-E) and found them to be broadly similar to those observed with IB-Ot infection (Fig. 3).

Together these data show that the general trend of EB infection is similar to that with IB, although the growth is slower and the timing of initiation is delayed.

## Discussion

In this study, we define the developmental progression of Ot during intracellular infection and reveal substantial phenotypic heterogeneity among individual bacterial cells. Through quantitative microscopy, we delineate four distinct intracellular growth stages—pre-active, lag, log, and maturation—each characterized by unique patterns of metabolic activity, DNA replication, and subcellular localization. We also demonstrate that even within a single microcolony, individual bacteria differ in the expression of key transcripts and proteins, revealing a high degree of asynchronous behavior and suggesting that Ot populations within host cells are developmentally diverse rather than uniform.

Our analysis of the early infection stages shows that a subset of Ot cells initiates protein synthesis within 1.5 hours of entry, while others remain metabolically quiescent. This variability implies that entry into the host cytoplasm does not synchronously activate bacterial metabolism. The presence of a defined lag phase, followed by a clear onset of DNA replication and cell division, supports the idea that Ot undergoes a structured developmental program within host cells, comparable to growth-phase transitions observed in free-living bacteria. The identification of a distinct maturation phase - marked by peripheral localization and reduced protein synthesis - suggests an adaptation for egress and survival outside the host cell, consistent with the previously described extracellular form of Ot that exhibits altered morphology and reduced metabolic activity^8^.

A key finding of this work is that the timing of the transition to maturation depends on the intracellular bacterial load rather than on chronological time post-infection. Cells harboring larger bacterial populations progress to the maturation stage earlier than those with fewer bacteria. This observation suggests that population density - or local nutrient limitation within the host cytoplasm - may serve as a cue for developmental progression. Such density-dependent regulation has parallels in other intracellular pathogens and may reflect a strategy to optimize the timing of host cell exit and reinfection.

Our single-cell analyses reveal striking transcriptional and translational heterogeneity within individual bacterial microcolonies. While genes involved in transcriptional regulation (e.g., *ctrA*, *trcR*, *rpoD*) and division (*ftsZ*, *ftsQ*) display distinct temporal expression patterns, we also observe spatial organization within colonies: regulators such as CtrA and TrcR are enriched in centrally located bacteria, whereas surface proteins such as ScaA and ScaC are more abundant at the periphery. This spatial differentiation suggests the presence of functionally specialized bacterial subpopulations, potentially reflecting a gradient of developmental states from metabolically active, dividing cells in the colony core to pre-egress, maturation-stage cells at the edge. Similar spatial heterogeneity has been described in biofilms and intracellular microcolonies of *Salmonella* and *Chlamydia*, where distinct metabolic zones and functional specialization support persistence and transmission^23,31^.

The regulatory hierarchy inferred from these data supports a model in which CtrA functions as a master regulator of developmental transitions in Ot, as it does in the related free-living Alphaproteobacterium *Caulobacter crescentus*^26,27^. CtrA expression peaks early, during the lag phase, preceding the upregulation of its putative target *trcR*, which in *Caulobacter* acts as a transcriptional regulator involved in the control of cell cycle–dependent gene expression^32^. The sequential induction of *ctrA* and *trcR* we observe is consistent with a conserved regulatory cascade controlling the transition from dormancy to replication. In this framework, the sporadic expression of *spoT*, encoding a (p)ppGpp synthetase/hydrolase, could mark a subset of cells entering a stringent-like response, priming them for growth arrest and differentiation into maturation-stage or extracellular forms. These findings provide a first glimpse of how a conserved alphaproteobacterial regulatory logic may have been adapted to the obligate intracellular lifestyle of Ot.

Our results underscore the necessity of single-cell approaches for studying obligate intracellular bacteria. Population-averaged assays such as bulk RNA-seq or qPCR obscure the underlying heterogeneity revealed here. Within a single infected cell, we observe coexisting bacterial subpopulations representing distinct physiological states—replicative, transitional, and pre-egress. Such heterogeneity likely contributes to the resilience of Ot infections, allowing subsets of bacteria to persist or escape host defenses while others continue replication. The co-existence of intracellular and extracellular developmental forms further complicates this picture, suggesting that Ot’s life cycle is both temporally and spatially distributed.

Finally, by comparing infections initiated with intracellular (IB) and extracellular (EB) forms, we show that both follow broadly similar developmental trajectories, although EB infections are delayed and exhibit slower growth. This delay likely reflects the need for EB to reactivate metabolic and replicative machinery upon host entry, analogous to the resuscitation of dormant bacterial forms in other systems. The conservation of the same four developmental stages in both infection routes suggests that Ot employs a robust, cyclical developmental program independent of its entry state.

In summary, this work provides the first detailed temporal and spatial framework for the intracellular developmental cycle of Ot. By integrating measures of metabolic activity, replication, and gene expression at the single-cell level, we reveal a complex, heterogeneous population structure that underpins the bacterium’s intracellular lifestyle. Future studies leveraging genetic perturbation and live-cell imaging will be critical to dissect the molecular mechanisms controlling transitions between these stages and to determine how phenotypic diversity contributes to persistence, virulence, and transmission in this neglected pathogen.

## Materials and Methods

### Cell lines and bacterial strains

This study used L929 mouse fibroblast cell line (ECACC Cat:85011425), human umbilical vein endothelial cells (Sigma Aldrich C-12203) and Ot strain Karp^33^.

### Mammalian and bacterial cell growth

L929 cells were cultured in DMEM (Gibco 11965092) supplemented with 10% FBS (Sigma-Aldrich F7524) and maintained at 37°C and 5% CO_2_ conditions. L929 cells were seeded at 6,000 cells per well of either 8-well µ-slide (ibidi 80826) or 8-well chamber slide (Lab-Tek^®^II 154534). Seeded L929 cells were allowed to grow for 2 days before infecting with Ot strain Karp at MOI 200:1 and then maintained at 35°C and 5% CO_2_ until the required time/day of fixation from 0 hpi (hour post infection) to 7 dpi (day post infection). For maintaining Karp stocks, purified *Ot* were stored in SPG solution (218 mM sucrose, 3.76 mM KH_2_PO_4_, 7.1 mM K_2_HPO_4_, and 4.9 mM potassium glutamate) at −80°C prior to use.

HUVEC cells were cultured in Human Large Vessel Endothelial Cell Basal Medium (Gibco M-200-500) supplemented with Large Vessel Endothelial Supplement (Gibco A1460801). HUVEC cells were seeded at 2000 cells per well in 8 well µ-slide (ibidi 80826) and allowed to grow overnight before infecting with Ot strain Karp at the desired MOI. Infected cells were maintained at 36°C and 5% CO2 until fixation.

### qPCR quantification

To quantify Ot, the bacteria were purified from various stages of growth. For extracellular bacteria (EB) isolation, the supernatant portion of the culture was transferred to a 50 ml centrifuge tube and centrifuged at 1,000 rpm, 4°C for 5 min to remove host cells. The supernatant was then transferred into a new 50 ml centrifuge tube and centrifuged at 20,000 xg, 4°C for 15 min to collect EB pellet. For intracellular bacteria (IB) isolation, old media in the culture flasks was replaced with 1.5 ml of fresh DMEM. Infected host cells on the adhering surface were then scrapped and transferred into 2 ml microcentrifuge tubes. The microcentrifuge tubes were then placed into Next Advance Bullet Blander^®^ (BBY24M-0710977) and homogenised at speed 8 for 2 min to break host cells and release IB from host cells. The cell suspensions were then centrifuged at 300 xg, for 3 min at room temperature to remove host cells. The supernatants were then transferred into new 1.5 ml microcentrifuge tubes and centrifuged at 20,000 xg, room temperature for 3 min to collect IB pellet. Hotshot DNA extraction was performed for EB and IB. EB and IB cell suspensions were collected for 50 µl by centrifuged at 20,000 xg. The pellets were resuspended in 25 µl lysis buffer solution (25 mM NaOH and 0.2 mM EDTA) and heated at 95°C for 30 min. The lysed solution is then added with 25 µl neutralization buffer (40 mM Tris-HCl) and stored at −20°C or directly used as a template in qPCR. qPCR was carried out according to protocols.io^34^. Amplification was performed on Bio-Rad CFX96 (CT008648) using qPCRBIO Probe Mix Lo-ROX (PCR Biosystems PB20.21), 47kDa probe ([6-FAM]-TTCCACATTGTGCTGCAGATCCTTC-[TAMRA]), 47kDa forward primer (TCCAGAATTAAATGAGAATTTAGGAC), and 47kDa reverse primer (TTAGTAATTACATCTCCAGGAGCAA). The quantification of Ot copy number was then determined relative to the 47kDa standard curve.

### CFSE quantification

To investigate bacterial proliferation rate over time, we performed Ot labelling with a stable protein binding fluorescent dye, carboxyfluorescein succinimidyl ester (CFSE). Purified Ot stocks were centrifuged at 20,000 xg for 3 min. The pellets were then resuspended in 1 ml phosphate buffered saline (PBS) (ThermoFisher 003002) supplemented with 1 µM CFSE (Ab113853) and incubated for 15 min in dark at room temperature. After the incubation, the mixture was added with 900 µl DMEM+10% FBS to quench dye for 5 min at room temperature. The labelled Ot is then centrifuged at 20,000 xg 15 min at 4°C. The pellet is then resuspended in DMEM+10% FBS and used to infect L929 seeded in 8-well µ-slide at MOI 1000:1. After 3 hours of incubation, live imaging was performed to visualize CFSE labelled Ot at 0 dpi. After the imaging, the cells were scraped from the 8-well µ-slide and performed DNA quantification by qPCR as mentioned above. These procedures were then repeated for 1-7 dpi time point.

### HPG labelling and quantification

To measure metabolic activity of the bacteria, we performed Click-iT HPG AlexaFluor protein Synthesis Assay (ThermoFisher C10269) at time points 0-6 hpi and also 0-7 dpi. On the day of fixation, the media was removed and replaced with minimal medium, DMEM lacking L-methionine (Gibco 21013024), supplemented with 40 µg/ml cycloheximide (Sigma-Aldrich C7698) and incubated for 30 min at 35°C. The media was then replaced with minimal medium supplemented with 50 µM L-homopropargylglycine (Click-iT^®^ HPG, C10269) and incubated for 1 hour at 35°C. The media were removed from the slides and then fixed with 4% formaldehyde for 10 min. Cell permeabilization was performed on ice by incubating in absolute EtOH for 1 hour followed by 0.5% Triton X-100 for 30 min. Blocking was performed at room temperature using PBS (pH7.4) containing 0.1% Tween 20 and 2% BSA for 30 min. The slides were then incubated with in-house-generated primary antibody (rat monoclonal antibody against TSA56 diluted 1:200) at 4°C overnight. The slides were then incubated with secondary antibodies goat anti-rat Alexa Fluor 488 (ThermoFisher A11006 diluted at 1:1000) supplemented with the nuclear stain Hoechst (1:2000 dilution) at 37°C for 1 hour. Click-iT reaction was then performed by incubating the cells with Click-iT cocktail (172 µl of 1X Click-iT reaction buffer, 8 µl of 100 µM CuSO_4_, 20 µl of 1X Click-iT additive, and 0.12 µl of 10 µM Alexa Fluor 594 azide (A10270)) at room temperature in dark. The washing procedure was performed with PBS (0.1% Tween 20, 2% BSA) each successive immunofluorescence labelling step. Samples were mounted using mounting media (20 mM Tris pH 8.0, 0.5% N-propyl-gallate, and 90% glycerol).

### Microscopy sample preparation Immunofluorescence

To measure the protein expression of CtrA, RpoD, TrcR, FtsQ, FtsZ, ScaA, and ScaC, Karp infected L929 cells were prepared for 0-7 dpi time course. On the required timepoint, the media were removed from the slides and then fixed with 4% formaldehyde for 10 min. Cell permeabilization was performed on ice by incubating in absolute EtOH for 1 hour followed by 0.5% Triton X-100 for 30 min. Blocking was performed at room temperature using PBS (pH7.4) containing 0.1% Tween 20 and 2% BSA for 30 min. The slides were then incubated with in-house-generated primary antibody diluted 1:200 (rat monoclonal antibody against TSA56) and target specific primary antibody diluted 1:200 (ABclonal rabbit monoclonal antibody against CtrA (WG-04957D), RpoD (WG-04962D), or TrcR (WG-04958D) or in-house-generated rabbit monoclonal antibody against FtsQ, FtsZ, ScaA, or ScaC) at 4°C overnight. The slides were then incubated with secondary antibodies—goat antirat Alexa Fluor 488 diluted at 1:1000 (ThermoFisher A11006) and goat antirabbit Alexa Fluor 594 diluted at 1:1000 (ThermoFisher A11037)—supplemented with the nuclear stain Hoechst (1:2000 dilution) at 37°C for 1 hour. The washing procedure was performed with PBS (0.1% Tween 20, 2% BSA) each successive immunofluorescence labelling step. Samples were mounted using mounting media (20 mM Tris pH 8.0, 0.5% N-propyl-gallate, and 90% glycerol).

### RNAscope

Integrated co-detection workflow (ICW) was followed according to RNAscope® Multiplex Fluorescent v2 Assay (ACD MK 51-150) to measure the RNA expression of *ctrA*, *rpoD*, *trcR*, *ftsQ*, *ftsZ*, *scaA*, *scaC*, and *spoT*. On the required timepoint (0-7 dpi), the media were removed from the slides and then fixed with 4% formaldehyde for 10 min. The cells were dehydrated with 50% EtOH, 70% EtOH, and 100% EtOH at room temperature for 5 min for each step. This is then followed by rehydration steps, where the slides were submerged in 70% EtOH, 50% EtOH, and 1X PBS. The slides were then treated with H_2_O_2_ (ACD322335) at room temperature for 10 min to block endogenous peroxidase. The slides were then washed with 1x Wash buffer (ACD310091) which was also used between each later step of RNAscope labelling. The slides were incubated with in-house-generated primary antibody diluted 1:20 (rat monoclonal antibody against TSA56) at 4°C overnight. Fixation with 4% formaldehyde was performed again to fixed the primary antibody for 30 min. Protease plus (ACD322331) diluted 1:15 was used to treat the samples at 40°C for 30 min to free RNA transcript from bound protein. Specific probes (*ctrA* (ACD1267591), *rpoD* (ACD1175811), *trcR* (ACD1267621), *ftsQ* (ACD1267601), *ftsZ* (ACD1267611), *scaA* (ACD1175821), *scaC* (ACD1169961), and *spoT* (ACD1176791)) were then hybridized using HybEZ^TM^ II Hybridization System (ACD 321711) for 2 hours at 40°C. The slides were stored in 5X saline sodium citrate (SSC) at room temperature overnight. Amplification steps were then performed by incubating the slides at 40°C using AMP1 (ACD323101) for 30 min, AMP2 (ACD323102) for 30 min, and AMP3 (ACD323103) for 15 min. Signal development was performed at 40°C by incubating the slides with HRP-C1 (ACD323104) for 15 min, Opal 570 (FP1488001KT) for 30 min, and HRP blocker (ACD323107) for 15 min. The slides were then incubated with secondary antibodies goat antirat Alexa Fluor 488 diluted at 1:1000 (ThermoFisher A11006) at room temperature for 1 hour. DAPI was used for the counter stain which was incubated in dark for 2 min at room temperature. Samples were mounted using ProLong^TM^ Gold Antifade Mountant (ThermoFisher P36930).

### Microscopy imaging

For Figures 1, 3 and 5, imaging was performed using a Zeiss Observer Z1 LSM700 confocal microscope with an HBO 100 illuminating system equipped with a Plan-APOCHROMAT 63x/1.4 oil objective lens (Carl Zeiss, Germany). Laser line 405 nm, 488 nm, 555 nm and phase contrast (T-PMT, DIC) were acquired using ZEN 2011 SP7 FP3 (version 14.0.0.0) software. All experimental images were captured under uniform imaging conditions, including fixed laser excitation, detector gain, and pinhole aperture settings.

For Figures 2 and 4, samples were imaged using a Nikon Ti2-E inverted microscope with a pE4000 Universal Fluorescence illumination system equipped with a 100XH Oil/1.45 CFI Plan-APOCHROMAT × _λ _objective lens (Nikon Europe B.V.) and LED-DAPI-A-2360A, LED-FITC-A-2745A, LED-TRITCA and LED-Cy5-5070A filter cubes (32mm). 3D deconvolution of Z-stacks was performed on Nikon NIS-Elements AR Version 5.42.04, followed by Maximum Intensity Projection.

### Microscopy analysis

Fluorescence intensity measurements and image segmentation for IFM and RNAscope were performed using automated pipeline using CellProfiler (4.2.7) software. For nucleus segmentation, diameter was set at 25-115 pixel units, threshold strategy set as adaptive-Otsu, threshold lower and upper bounds of 0.1-1.0, and clump distinguish using shape and propagate method. For TSA56 labelled *Ot* segmentation, diameter was set at 1-40 pixel units, threshold strategy set as adaptive-minimum cross-entropy, threshold lower and upper bounds of 0.05-1.0, and clump distinguish using intensity and propagate method. For specific target labelled *Ot* segmentation, diameter was set at 1-40 pixel units, threshold strategy set as adaptive-minimum cross-entropy, threshold lower and upper bounds of 0.03-1.0, and clump distinguish using intensity and propagate method. For low intensity specific target signal seen in RNAscope analysis, the threshold lower limit was changed to 0.01. For clump analysis, images obtained from 4-7 dpi were used with clump diameter set at 50-300 pixel units, threshold strategy set as adaptive-minimum cross-entropy, threshold lower and upper bounds of 0.01-1.0, and clump distinguish using shape and propagate method. For CFSE, the fluorescent intensity was measured manually using Fiji Image J software. The bacteria were quantified by measuring the brightest pixel intensity.

The statistical analyses for band intensity were carried out using RStudio (2025.05.1) and GraphPad Prism version 10.4.2 (633). A Mann-Whitney test was used to compare intensity between two sample groups. Kruskal-Wallis test was used to compared differences between 3 or more groups. Pearson’s correlation was used to test the relationship between intensity and *Ot* position. Symbol ns indicates no significant, * indicates p-value <0.05, ** indicates p-value <0.01, *** indicates p-value <0.001, and **** indicates p-value <0.0001.

Statistical analyses where response variables were proportional (Figure 2) were carried out using RStudio (2025.05.1). A generalised linear model (GLM) was fitted to assess the effect of Timepoint, MOI and their interaction on proportion of maturation cells. Nested models were compared using a likelihood ratio test via a sequential Analysis of Deviance; significance of each term was evaluated by chi-square tests based on the reduction in deviance when this term was added to the model.

---

**Supplementary Figure 1. CFSE growth curve**

Growth curve showing Ot growth, measured by genome copy number measured by qPCR with primers against the single copy gene *tsa47.* The bacterial levels in the presence of 1 µM CFSE and no CFSE are shown. The addition of 1 µM CFSE does not significantly decrease bacterial growth.

**Supplementary Figure 2. Immunofluorescence microscopy analysis of Ot during infection.**

Representative images of immunofluorescence microscopy of Ot infected in L929 cells fixed at 4-, 5- and 6-days post infection, showing localisation of bacteria expressing CtrA, FtsQ, FtsZ, RpoD, ScaA, ScaC and TrcR. Immunofluorescence microscopy labels proteins using a polyclonal antibody against the protein of interest in red. The bacteria are counterstained using an antibody against TSA56 in green and host nuclei are labelled with Hoechst in blue. Scale bar = 10 µm

**Supplementary Figure 3. RNAscope analysis of Ot during infection.**

Representative images of RNAscope microscopy of Ot infected in L929 cells fixed at 4-, 5- and 6-days post infection, showing localisation of bacteria expressing *CtrA, FtsQ, FtsZ, RpoD, ScaA, ScaC, SpoT* and *TrcR.* RNAscope labels bacterial transcripts using a probes against the gene of interest in red. The bacteria are counterstained using an antibody against TSA56 in green and host nuclei are labelled with Hoechst in blue. Scale bar = 10 µm

